# Different helminth parasites show contrasting relationships with age in a wild ungulate

**DOI:** 10.1101/2022.10.31.514008

**Authors:** Gregory F Albery, Sean Morris, Alison Morris, Fiona Kenyon, David McBean, Josephine M Pemberton, Daniel H Nussey, Josh A Firth

## Abstract

As animals age they often exhibit substantial physiological and behavioural changes that can drive changes in infection status over the lifespan. Generally, immunosenescence is expected to lead to greater infection in older individuals, but this process could be confounded or counteracted by changes in behaviour, selective disappearance of certain individuals, or a range of immune processes. Here, we uncover contrasting age-related patterns of infection across three different helminth parasites in wild adult red deer (*Cervus elaphus*). Counts of strongyle nematodes (order: Strongylida) increased with age, while counts of liver fluke (*Fasciola hepatica*) and tissue worm (*Elaphostrongylus cervi*) decreased. These relationships could not be explained by selective disappearance of certain individuals or changes in behaviour, suggesting that immune changes could be responsible. Additionally, we found a positive relationship between social connectedness and strongyle infection, implying that previously documented age-related decreases in social behaviour may minimise exposure, reducing the negative effects of immunosenescence. These findings demonstrate that burdens of different parasites can show contrasting changes over an individual’s lifespan depending on a complex suite of intrinsic and extrinsic factors.

## Introduction

An animal’s disease burden is dependent on a combination of its susceptibility and exposure to parasites [1,2], As an animal ages, it undergoes a suite of physiological changes [3], many of which affect the immune system [4–6], Because these changes often result in increased susceptibility to infection, it is usually expected that individuals will exhibit a greater prevalence or burden of parasites as they age [6–8]. However, this process is potentially confounded by a number of age-related processes: first, the immune system can change and adapt in many other ways over the lifespan; most importantly, individuals acquire adaptive immunity to certain parasites as they become infected, potentially leading to an increase in immunity to these particular parasites [9]. Second, individuals alter their behaviour as they age [10,11], which could alter their exposure rate and therefore their infection probability; for example, individuals could alter their feeding locations [12], or they could become more or less social [10,13], both of which could expose them to different parasites or at different rates. Third, because parasites often exact survival costs on their hosts, more heavily-infected individuals may be more likely to die – a process known as “selective disappearance” – which could produce a negative age-infection trend at the population level, and may bias estimates of within-individual ageing patterns [14,15]. The emergent pattern of infection status over the lifespan will depend on a combination of these factors.

Given these multiple processes, ageing individuals could generally show greater burdens of all parasites, or an individual’s infection statuses could be asynchronous and divergent, leading to an age-related shift in parasite community composition. For example, individuals may move from one area to another as they age [13] and therefore become exposed to a different variety of environmental parasites. Alternatively, due to molecular differences between parasites and differing immune evasion capacities, individuals may gain immunity to certain parasites (such that older individuals are less often infected) while remaining unable to develop an effective immune response to others (such that age has no relationship with infection overall).

As yet, it is unclear how these processes combine to influence age-related changes in infection in wild animals. Despite some documented examples of age-infection relationships [e.g. [16–21]], most previous studies are cross-sectional rather than longitudinal, and are therefore unable to identify and extricate selective disappearance effects [[14,15]; but see [16,17,20]]. Further, most studies focus on one parasite, and it therefore is unclear how often parasites show divergent age-related trends within a population. Here, we address this gap by asking how different helminth parasite counts change over the lifespan in a long-term study population of wild red deer (*Cervus elaphus*). By examining changes in socio-spatial behaviour and relationships between counts of multiple parasite taxa and survival, we assess whether these changes occur within individuals, and how they might operate through exposure-related pathways.

## Methods

### Study population

The study population was the individually-monitored Isle of Rum red deer; this unmanaged wild population has been studied since 1973 [22], with regular faecal parasite sampling since 2016 [23]. The deer are censused 40 times a year, with individuals known by name and tagged using a combination of methods. The deer give birth in May and June, and daily censuses over the calving period allow >90% of calves to be caught, tagged, and weighed. The deer year runs from 1^st^ of May, and individuals are assigned an age in years based on the year they were born in; for example, all individuals turn 1 year old on the 1^st^ of May the year after they were born. 40 study area censuses per year allow us to keep track of each individual’s life history, and individuals have known death dates, generally to within one month, and often to the day, allowing accurate quantification of mortality effects. Following our previous related work in this system [13,24], here we assess mature females (3 years and older), as these are the best-understood age and sex class, with the largest available dataset; young males disperse and few adult males live in the study area and so male are less well sampled. Female reproductive status in any year was coded as either “none” (did not give birth that year), “summer” (lost calf before October 1^st^), or “winter” (lost calf over its first winter or reared it through its first winter).

### Parasitology

We have previously described our parasitology monitoring regime in detail [23]. Briefly, three times a year (late April, August, and November), for two weeks at a time, we observe the deer intensively to collect faecal samples from as many individuals as possible. After observing an individual defaecating, we collect the sample as soon as possible into a resealable plastic bag, and at the end of the day we homogenise it, and store it anaerobically in the fridge (~4°C).

We counted gastrointestinal helminth parasite propagules within three weeks. We counted strongyle nematode (order: Strongylida) eggs using a salt flotation-centrifugation technique [23]; liver fluke (*Fasciola hepatica*) eggs using a sedimentation technique; and tissue worm (*Elaphostrongylus cervi*) and lungworms (*Dictyocaulus* sp.) larvae using a Baermannisation technique. All techniques are accurate to at least 1 egg or larva per gram. Our salt flotation also detected a number of other parasites (described in [23]), but they were present at low prevalence (<10%) in adult females, and therefore we were less able to analyse how they changed with age.

Samples were collected between August 2016 and April 2021. Where multiple samples were collected for a given individual in a given sampling trip, we took the mean of the counts to leave a maximum of one count per individual per sampling trip. Our final dataset included Ns=1439 measurements taken from Ni=210 individuals; some assays were not completed for all samples, leaving Ns=l433 *F. hepatica* measurements, and Ns=1126 *E.cervi* and *Dictyocaulus* measurements taken from Ni=209 individuals.

### Behavioural pipeline

We constructed social networks as previously described [13,24], using all census observations of each individual in each year, including adults and juveniles. We chose to include juveniles in the social network as they are heavily infected with parasites [23] and could therefore play an important role in infecting older individuals. Social connections were judged by field workers based on a spatially-parameterised “gambit of the group” approach, where individuals within a certain distance were taken to be socialising (see [13,24] for details). We also calculated local density as previously described [13,24], using all observations of each individual in each year, including both adults and juvenile individuals. This approach uses a kernel density estimator, taking individuals’ annual centroids and fitting a two-dimensional smooth to the distribution of the data. Individuals are then assigned a local density value based on their annual location on this kernel.

We examined how an individual’s social connectedness the previous year was associated with its parasite burden in the focal year. Using the previous year’s social network allowed us to accommodate the time lag of the influence of social connections on parasite burden (e.g. including parasites’ time to development and maturation and egg production) and also allowed us to avoid confounding produced by analysing an individual’s social connectedness in a given year with its concurrent and earlier parasite infection status, or reverse causality emerging from e.g. avoidance responses [25]. As such, we examined how an individual’s contacts from May 1^st^ in year t to April 30^th^ in year t+1 affected its parasite burden in August year t+1, November year t+1, and April in year t+2. Although a relatively coarse annual measure of sociality, individual-level repeatability of social network location is high [24], as is spatial fidelity [26,27], and previous work has shown these measures to be ecologically relevant for individuals [13,24], As such, we judge this to be a reliable indicator of social connectedness with relevance to the risk of parasite transmission over the lifespan.

### Models

Our dataset included 1449 seasonal measures of 210 individual deer, spread across 5 deer years. To identify age-related changes in parasite burden and determine their causes, we fitted a selection of generalised linear mixed models (GLMMs) using the Integrated Nested Laplace Approximation (INLA) in R [28]. INLA is a deterministic Bayesian algorithm that allows fitting of spatially distributed random effects to account for spatial autocorrelation in the response variable [29]. The model sets we used were as follows:

**Base models:** first, we examined each parasite count as a response variable with a negative binomial distribution. We fitted explanatory variables including Year (factor with 5 levels: 2016-2020); Season (factor with 3 levels: Summer, Autumn, and Spring); Reproductive Status (Factor with 3 levels: None, Summer, and Winter); Age (continuous covariate, range 3-24, mean 7.9); Degree Centrality (continuous covariate, range 0-290, mean 115.4); and Local Density (continuous covariate, rescaled to have values between 0 and 1). We ran these models both without and with a random effect of individual identity, to examine how controlling for among-individual variation impacted our estimates of age effects.
**Spatial models:** second, to identify whether our results were affected by spatial autocorrelation, we added a spatially distributed Stochastic Partial Differentiation Equation (SPDE) effect [29–31] to our base parasite models. Fitting this effect had several purposes: by comparing the fit of the spatial model with the base model, we could identify whether the parasite counts were significantly spatially autocorrelated; by comparing the model estimates we could identify whether this spatial autocorrelation was affecting our conclusions; and by plotting the effect in space we could identify spatial hot- and coldspots of infection [31]. To assess model fit, we used deviance information criterion (DIC), with a cutoff of −2 ΔDIC to distinguish between competitive models.
**Survival models:** often, ageing models incorporate fixed effects of longevity to examine selective disappearance of certain individuals [14]. We were unable to do this with our dataset, as it spanned five years running to the present; because many individuals were yet to die, we did not have known longevity values for many of the data points, which reduced our models’ power in this context. To provide an approximate answer to this question, we fitted binomial survival models following previous methodology [32] to examine whether parasites were likely to be causing mortality. Overwinter survival (0/1) was fitted as a response variable, with explanatory variables including Year; Reproductive Status; Age; Degree Centrality; Local Density; and a random effect of individual identity, all as described above. We sequentially added each parasite count (log(X+1)-transformed) as an explanatory variable, one at a time, to investigate whether they correlated with subsequent survival. We note that this is a relatively crude way of assessing selective disappearance effects that was necessitated by our dataset; depending on the effects shown by the mortality assessments, we may or may not be able to infer an effect of selective disappearance using such an analysis.

## Results

All effect estimates and credibility intervals are given in Supplementary Figure 1-Y. We found substantial age effects on counts of three parasites: there were positive associations between age and strongyle count, and negative associations between age and liver fluke and tissue worm count (Figure 1A-C; P<0.001 for all three). Lungworms, meanwhile, showed no relationship with age (Figure 1D; P>0.05).

**Figure 1:**
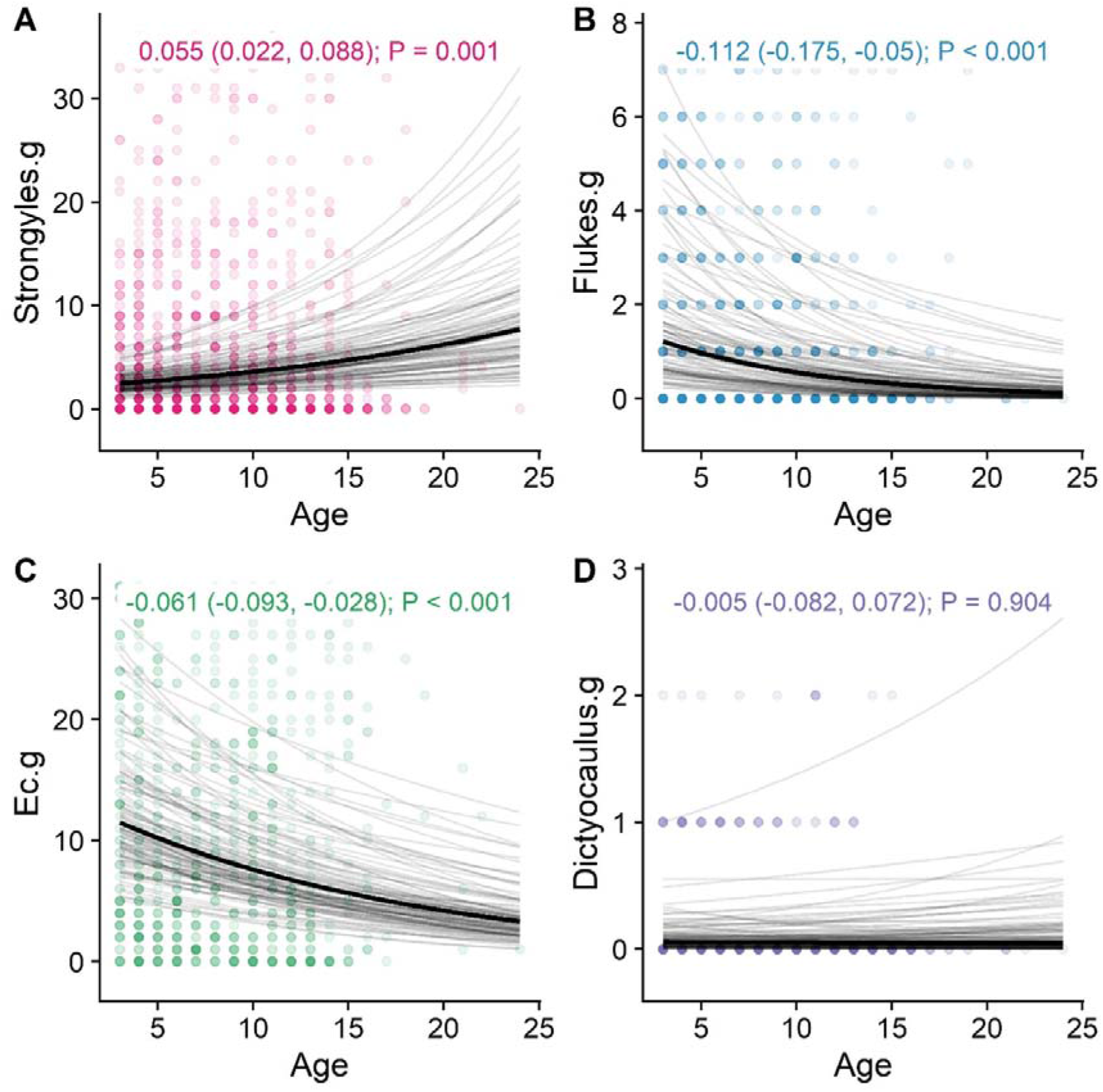
Age-related changes in infection with four helminth parasites in wild red deer. A, strongyle eggs per g; B, Fluke eggs per g; Cy, *E. cervi*/larvae per g; D, *Dictyocualus* larvae per g. Taken from the best-fitting models, the dark black line represents the mean of the posterior distribution for the age effect estimate; the light grey lines are 100 random draws from the posterior to represent uncertainty. The age effect estimate, credibility intervals, and P values are given at the top of each panel. The points represent individual samples, with transparency to allow for visualisation of overplotting. The figures have been cropped to the distribution of the fitted lines to help visualising the model fits, so some extreme points have been excluded from the figures.

We also uncovered a moderate positive effect of social connectedness on strongyle infection (Figure 2; 0.004, 0.001-0.006, P=0.012), but not in any other parasite model (Supplementary Figure 1; P>0.05).

**Figure 2:**
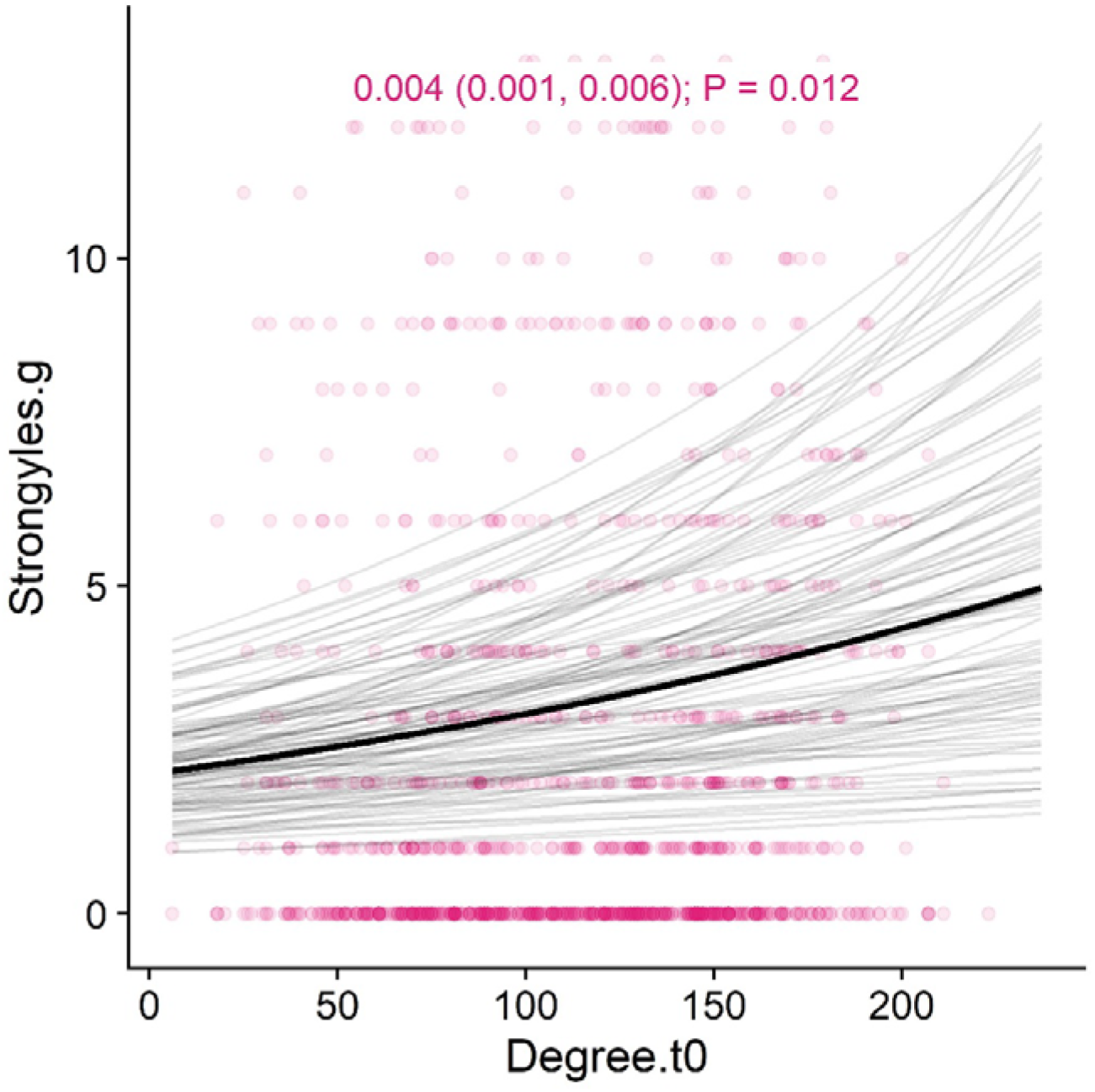
Association between social connectedness (degree centrality) in the previous year and strongyle nematode count in wild red deer. The x axis is in numbers of contacts; the y axis is in eggs per gram. Taken from the spatial model, the dark black line represents the mean of the posterior distribution for the age effect estimate; the light grey lines are 100 random draws from the posterior to represent uncertainty. The degree effect estimate, credibility intervals, and P values are given at the top of the figure. The points represent individual samples, with transparency to allow for visualisation of overplotting. The figure has been cropped to the distribution of the fitted lines to help visualising the model fits, so some points have been excluded from the figure.

Spatial autocorrelation effects substantially improved the models for flukes and tissue worms (Supplementary Table 1; ΔDIC<-3), but not for strongyles or lungworms (Supplementary Table 1; ΔDIC>-2). These findings demonstrate that there was notable heterogeneity in parasite infection (Supplementary Figure 3), but controlling for this effect did not impact our age estimates (Figure 3A, Supplementary Figure 1), demonstrating that changes in spatial location were unlikely to be responsible for our observed age effects. There were moderate density effects evident in the base models for *E. cervi* and *F. hepatica*, but these effects were removed when spatial autocorrelation was controlled for (Supplementary Figure 1). The spatial distributions of these parasites largely agreed with earlier observations [31], with greater *F. hepatica* count in the south-middle of the study area and greater *E. cervi* count in a slow gradient moving towards the north, particularly the northeast (Supplementary Figure 3).

**Figure 3:**
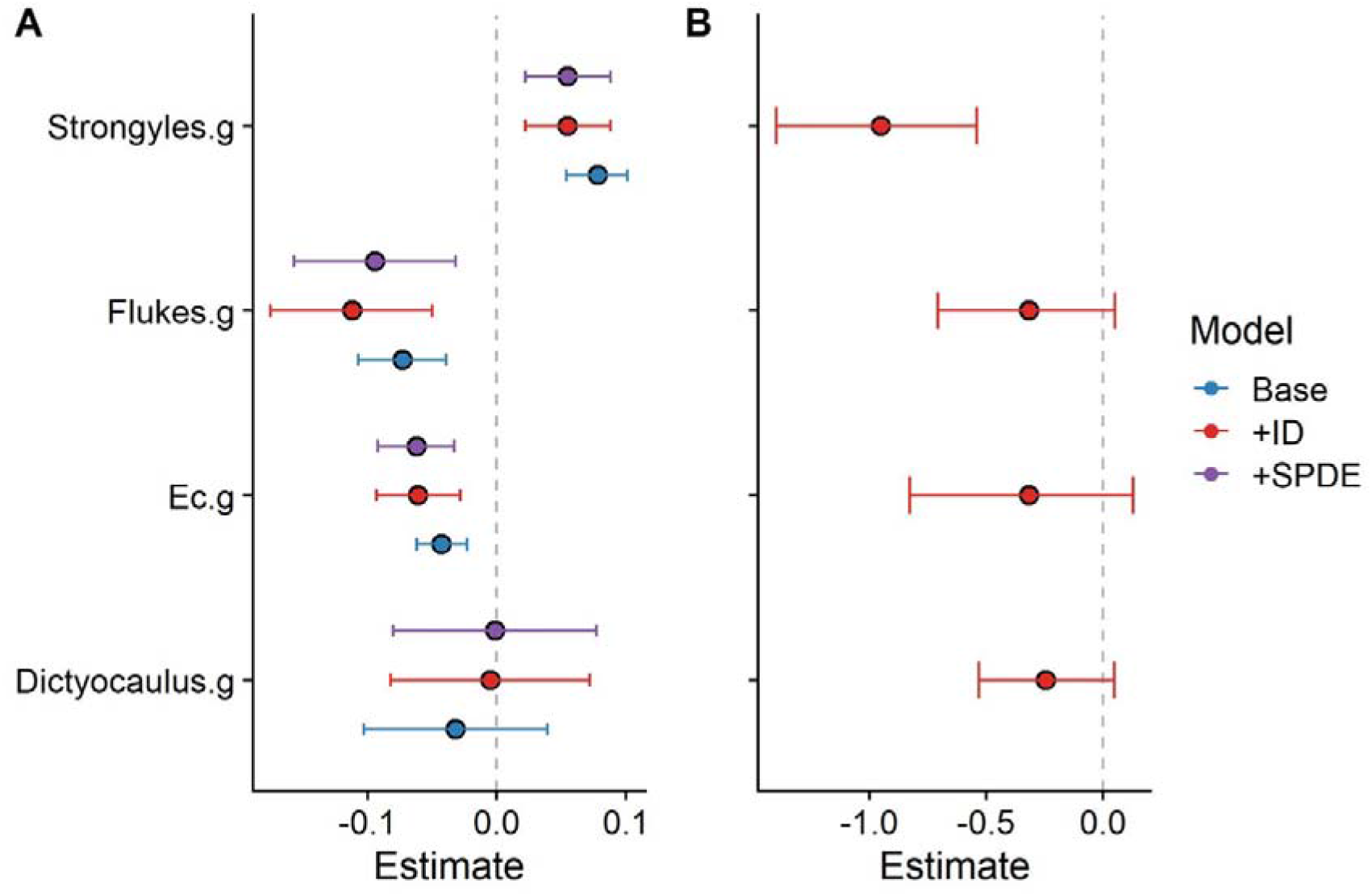
Model effect estimates for A) the effect of age on parasite counts and B) the effect of parasite counts on overwinter survival probability. Points represent the mean for each effect estimate; error bars denote 95% credibility intervals. All estimates are given on the link scale.

We found strong associations between strongyle count and overwinter survival probability (Figure 3B; Supplementary Figure 2; −0.98, −1.47- -0.55, P<0.001), agreeing with previous findings [32], There were only weak negative nonsignificant trends with the other parasites (Figure 3B; Supplementary Figure 2; P>0.05). Additionally, fitting random effects of individual identity substantially improved model fit (ΔDIC<-10) but without notably affecting the age effect estimates (Figure 3A; Supplementary Table 1). Taken together, these findings provide little evidence for a role of selective disappearance in *driving* our observations, except for potentially *obscuring* the age-strongyle trend. That is, our estimate for the age effect on the strongyle counts is a composite that likely includes a contrasting effect of selective disappearance, and is therefore likely an underestimate.

## Discussion

We uncovered substantial and contrasting changes in infection burden across different parasites over the lifespan in a long-lived wild mammal. As would be expected given a waning immune system, we indeed observed increased strongyle nematode counts with age; this observation adds to a relatively sparse body of evidence for age-related increases in parasite count in wild animals, particularly in longitudinal contexts [6,7,16,17,20,33]. However, interestingly, we also found concurrent decreases in liver fluke (*Fasciola hepatica)* and tissue worm (*Elaphostrongylus cervi*) counts. Given the limited influence of spatial autocorrelation or direct socio-spatial behavioural metrics in our models, these findings suggest that these changes were not driven by previously documented age-related movements to certain geographic areas or changes in contact rates [13]. Similarly, there was no evidence that selective disappearance of certain individuals was driving our observed trends, given limited survival effects of infection. Our dataset was limited by unknown death dates of many individuals, because they are still alive, necessitating the use of survival models as a proxy for selective disappearance. Nevertheless, the positive age effect in strongyles would only be blunted by the observed mortality effect; further, given the strength of the age effects in *F. hepatica* and *E. cervi* infection, extremely strong mortality would be required to drive our observations. As such, we are confident that the age-related changes more likely occurred due to intrinsic changes occurring over the course of individuals’ lives.

A variety of immune- and behaviour-related changes could be responsible for divergent age trends among parasite taxa: on the immune side, increasing strongyle counts could be occurring due to immunosenescence causing decreased resistance, agreeing with previous observations in wild Soay sheep [7,34], This observation disagrees with a previous finding that strongylid infection decreases with age in African elephants at the population level [21]; it is possible that selective disappearance may have played a role in influencing this pattern in the elephants. Meanwhile, the decreasing liver fluke and tissue worm counts could be indicative of acquired immunity over the lifespan. It is unclear how and why these trends would diverge for these two groups of parasites; confirming a role for the immune system would require 1) measuring a suite of immune traits to examine how they change with age, and 2) examining whether they correlate with parasites and could therefore represent immune resistance [35]. Given that the strongyle counts occur at the order level, and generally comprise a mixture of different species, one possibility is that even within this parasite count there is age-related change in the community, with certain species dominating in early years that are then replaced by higher-intensity infections with other species. Confirming this would require more precise taxonomic identification of the constituent nematodes, e.g. through DNA-based approaches [36,37], A similar trend is less likely for the fluke and tissue worm counts, as these are more likely to be counts of single homogenous species. Ultimately, the fact that these reputedly-similar macroparasites showed highly divergent trends is interesting, and invites further investigation.

Although our examined behavioural measures could not explain the observed changes in parasite count, it is likewise possible that other age-related changes in behaviour could play a role in the age-infection relationships. For example, animals often alter foraging behaviour as they age [11] and use social information to inform what they eat [38,39]. Given that older individuals may often be better informed, particularly in long-lived species [10], it is possible that older deer gradually learn – either by trial and error or from conspecifics – which forage areas or species are more likely to be contaminated with *F. hepatica* and *E. cervi*. In that case, older individuals would become gradually less infected with these parasites, creating a negative age trend without necessitating any immune changes. At the population level, such age-dependent habitat selection processes could drive the observed spatial population structuring [13] and produce an age-dependent “landscape of disgust” [33,40]. Both *F. hepatica* and *E. cervi* are transmitted through intermediate snail hosts, while strongyles transmit directly; in particular, *E. cervi* infection requires ingestion of the snail directly, and *F. hepatica* encysts on waterlogged vegetation which is then consumed. It is possible that both of these life stages are easier for older deer to learn to associate with illness and avoid, while directly ingested strongyle larvae do not afford this possibility.

Our observation that greater social connectedness was predictive of greater parasite count agrees with the conventional wisdom that infectious disease is a primary cost of sociality [41,42], but this trend was in the opposite direction to the direction we expected if social behaviour was playing a role in driving age-infection relationships. That is, if individuals’ ageing behaviour were driving the effect, because social connectedness decreases with age [13], we would expect social connectedness and parasite count to have a negative relationship rather than a positive one. Instead, these findings are more suggestive of the reverse: ageing individuals may be decreasing their social connectedness to reduce their exposure to parasites, potentially minimising the effects of a waning immune system [10,13]. The strength of selection often wanes in later life [43,44], and therefore it is unlikely that this is an adaptive response specifically brought on by immunosenescence; instead, this effect could emerge through more general behavioural compensation for a weak immune response that evolved in earlier life and persists as the animal senesces. Such behavioural compensation is relatively common [45,46]: for example, Stephenson [47] demonstrated that guppies (*PoeciHa reticulata*) show stronger conspecific avoidance when they are more susceptible to infection. Although it has yet to be shown that immunosenescence and social ageing are linked directly, our observations are consistent with a similar underlying process.

Connectedness in the social network was a better predictor of strongyle count than was a spatial measure of local density. This was unexpected, as given that helminth parasites transmit indirectly, we would expect that incorporating spatial measures (rather than more direct measures of social contact) would be more representative of indirect contact rates – and therefore of parasite counts [12]. There are several possible explanations for this: first, because social connections are temporally parameterised (i.e., they require individuals to be in the same location at the same time), the measures derived from this metric could be more indicative of helminth transmission, which could occur more on the timescale of weeks to months than years; second, individuals’ social connectedness may actually be a better indicator of their experienced local density than the annually summarised population density measures that we used; third, because social connections are estimated using spatial proximity in this system – rather than e.g. direct contact events like grooming – social connectedness may incorporate a substantial amount of information concerning spatial connectedness on the timescales relevant to helminth transmission. All these findings agree with the previous observation that social network position is both heavily intertwined with spatial behaviour in this system and a biologically important stand-alone measure [13,24]. Regardless of the ultimate cause, this finding adds notably to the literature on spatial-social analysis in disease ecology, and accentuates the value of using both spatial and social metrics when quantifying the drivers of infection status [12].

Overall, our results confirm that age-related changes in infection can vary substantially within the same system, and likely depend on a complex combination of immune, behavioural, and demographic processes. Ageing individuals may experience not only a greater overall parasite burden, but also a different parasite community, which may exert complex pressures on the age structure of the population. Understanding how and why parasite community structure changes with host age is likely to provide new insight into disease transmission and the ageing process in natural systems.

## Acknowledgements

We thank NatureScot and its predecessors for permission to work on the Isle of Rum NNR. The field project has been supported by grants mainly from the UK NERC with some additional funding from BBSRC, the Royal Society and ERC. We thank all who have contributed to the maintenance of the project over time, especially Loeske Kruuk. We thank multiple dedicated field workers who have contributed to field data collection, especially Fiona Guinness who collected the first 20 years of census data. GFA was funded by NSF grant number 1414296, and by a Bruce McEwen Career Development Fellowship the Animal Models for the Social Dimensions of Health and Aging Research Network (NIH/NIH R24 AG065172). JAF was supported by funding from BBSRC (BB/S009752/1) and NERC (NE/S010335/1 & NE/V013483/1).

## Supplementary Information

**Supplementary Table 1:**
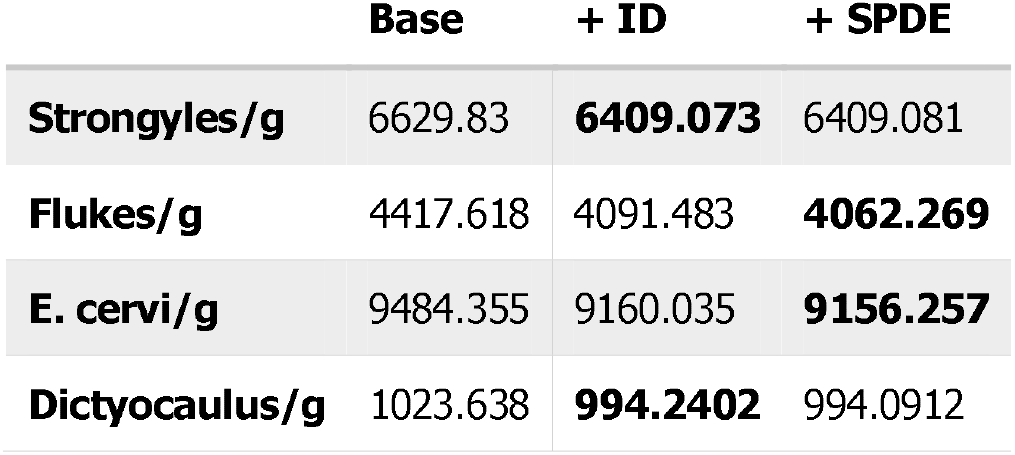
DIC values associated with the addition of individual identity effect f’+ID”) and spatially autocorrelated random effects (“+SPDE”). Values in bold represent the best-fitting model for each parasite.

|  | Base | + ID | + SPDE |
| --- | --- | --- | --- |
| Strongyles/g | 6629.83 | **6409.073** | 6409.081 |
| Flukes/g | 4417.618 | 4091.483 | **4062.269** |
| E. cervi/g | 9484.355 | 9160.035 | **9156.257** |
| Dictyocaulus/g | 1023.638 | **994.2402** | 994.0912 |

**Supplementary Figure 1:**
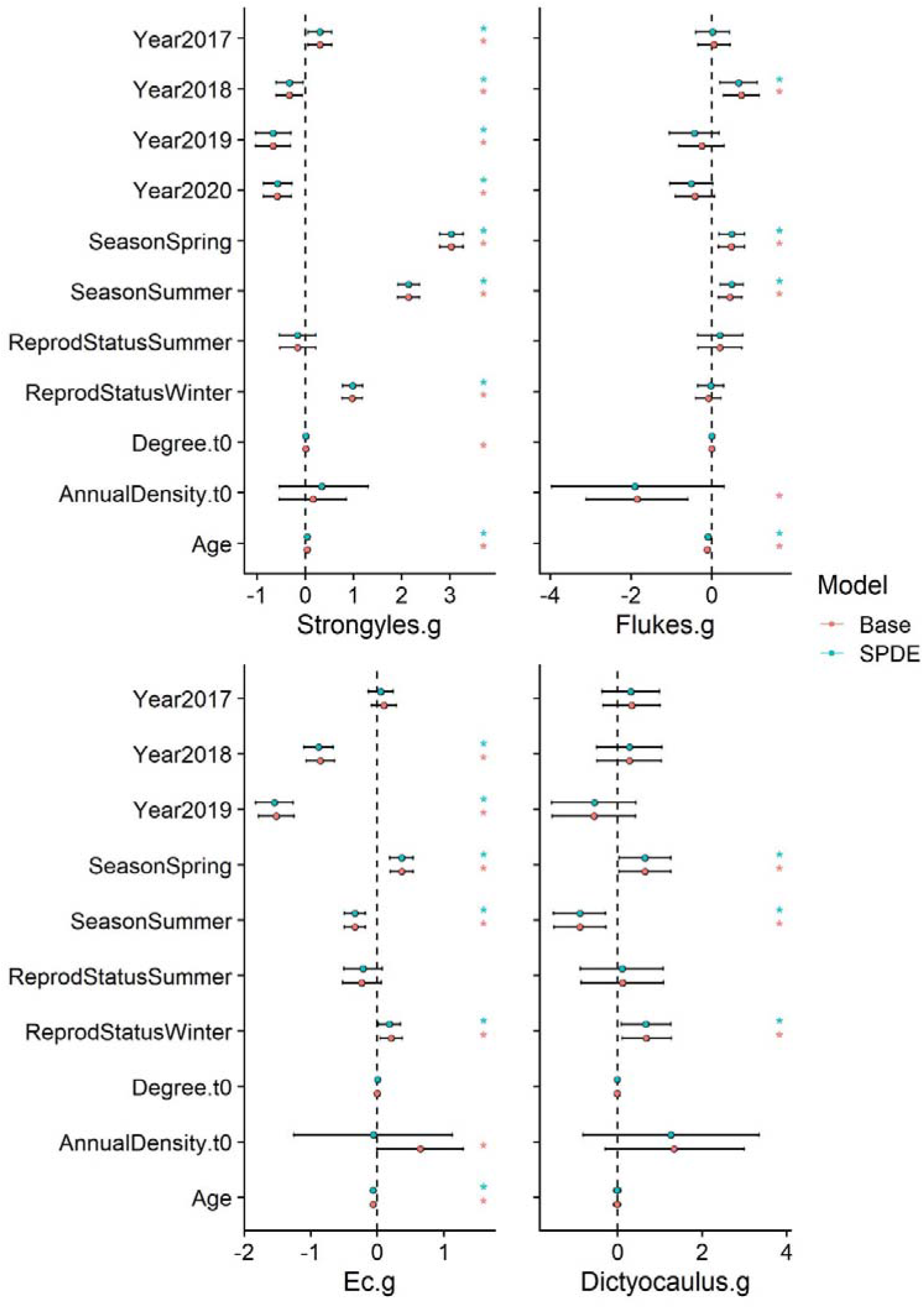
Model effect estimates for the full parasite count models, both without and with spatial autocorrelation effects. Points represent the mean for each effect estimate; error bars denote 95% credibility intervals. All estimates are given on the link scale. Estimates marked with an asterisk were significant (i.e., their credibility intervals did not overlap with zero).

**Supplementary Figure 2:**
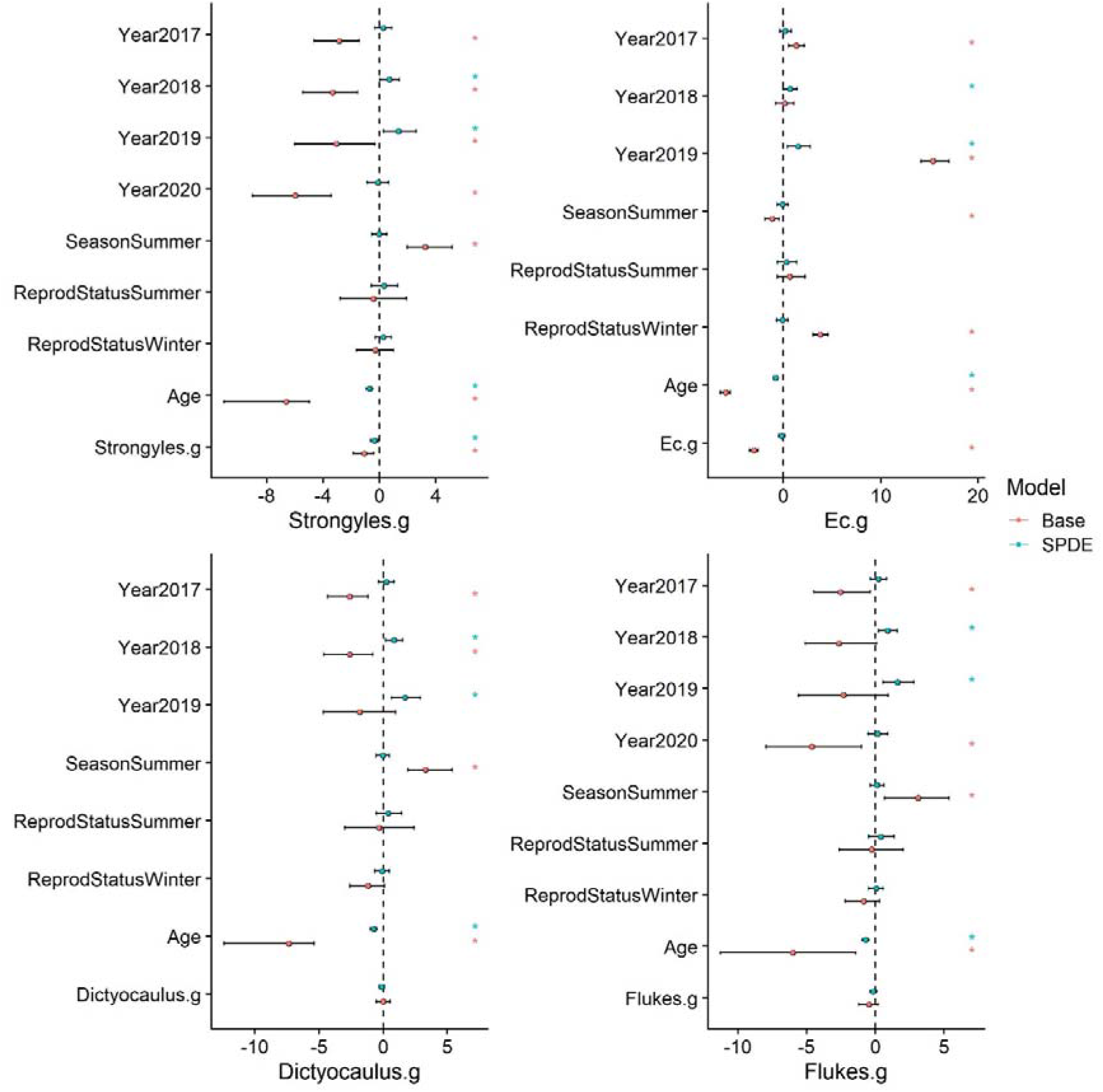
Model effect estimates for the survival models, both without and with spatial autocorrelation effects. Points represent the mean for each effect estimate; error bars denote 95% credibility intervals. All estimates are given on the link scale. Estimates marked with an asterisk were significant (i.e., their credibility intervals did not overlap with zero).

**Supplementary Figure 3:**
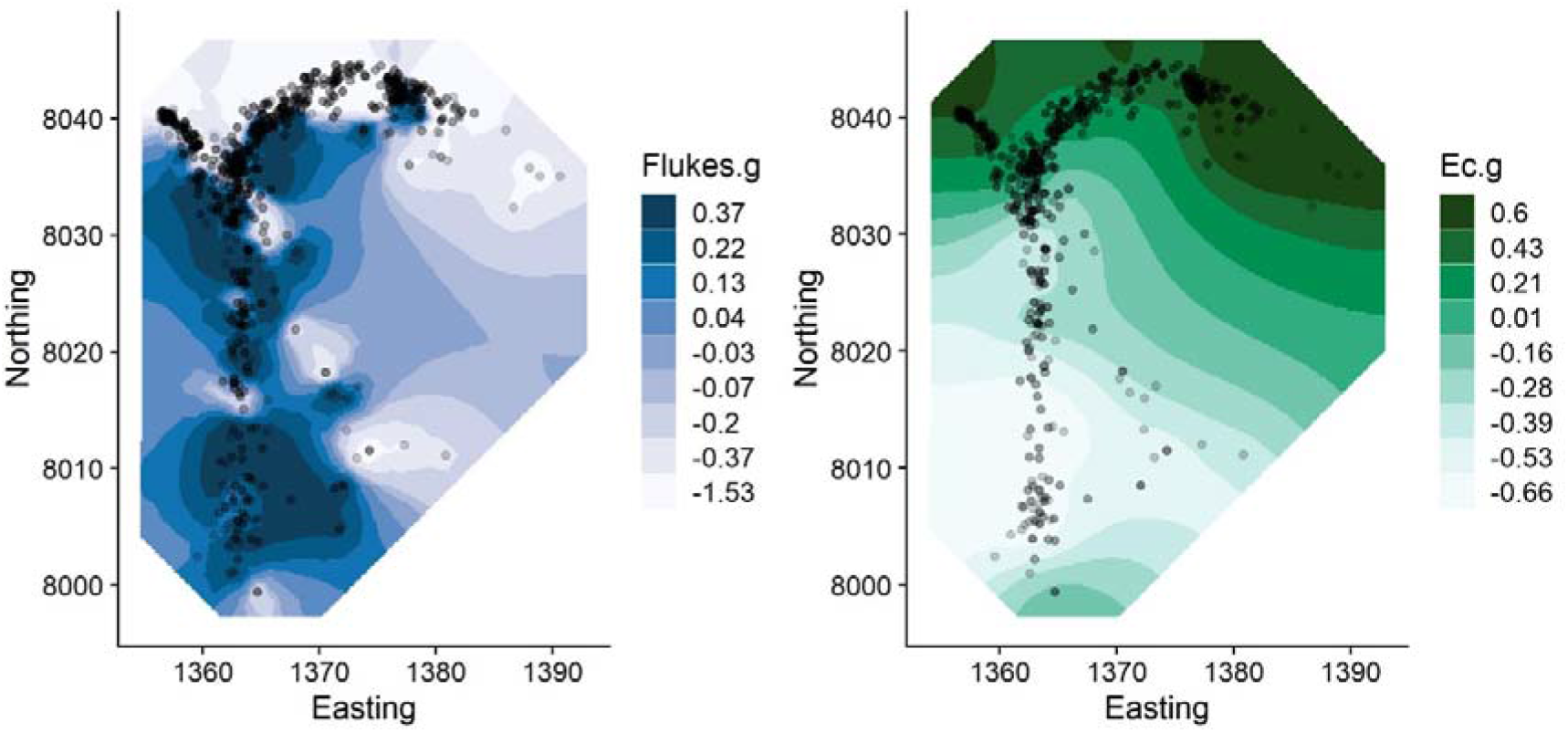
The distribution of spatial random effects (SPDE) for *E hepatica* and *E. cervi*, the only two significantly spatially heterogeneous parasites we examined. Darker colours represent greater parasite counts.

